## Supplementary figures and images for "Substrate-Specific Function of PNPLA3 Facilitates Hepatic VLDL-Triglyceride Secretion During Stimulated Lipogenesis"

### Supplemental Figure S1

#### Supplemental Figure S1

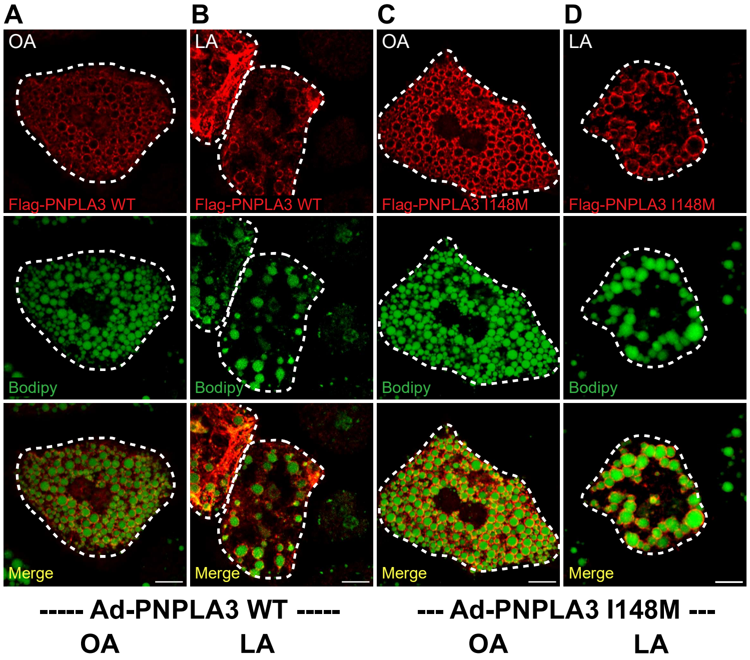

### Supplemental Figure S2

#### Supplemental Figure S2

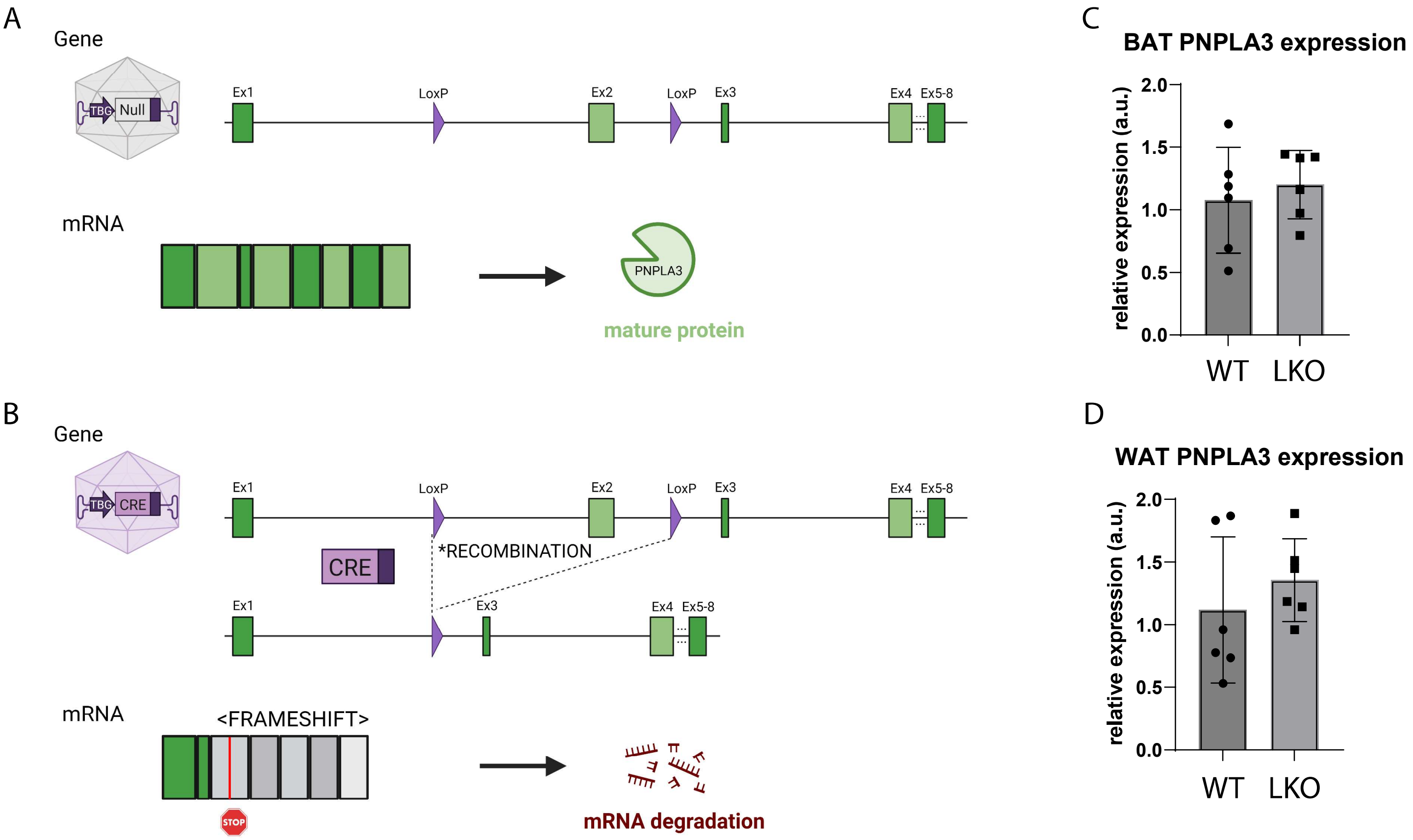

### Supplemental Figure S3

Supplemental Figure S3

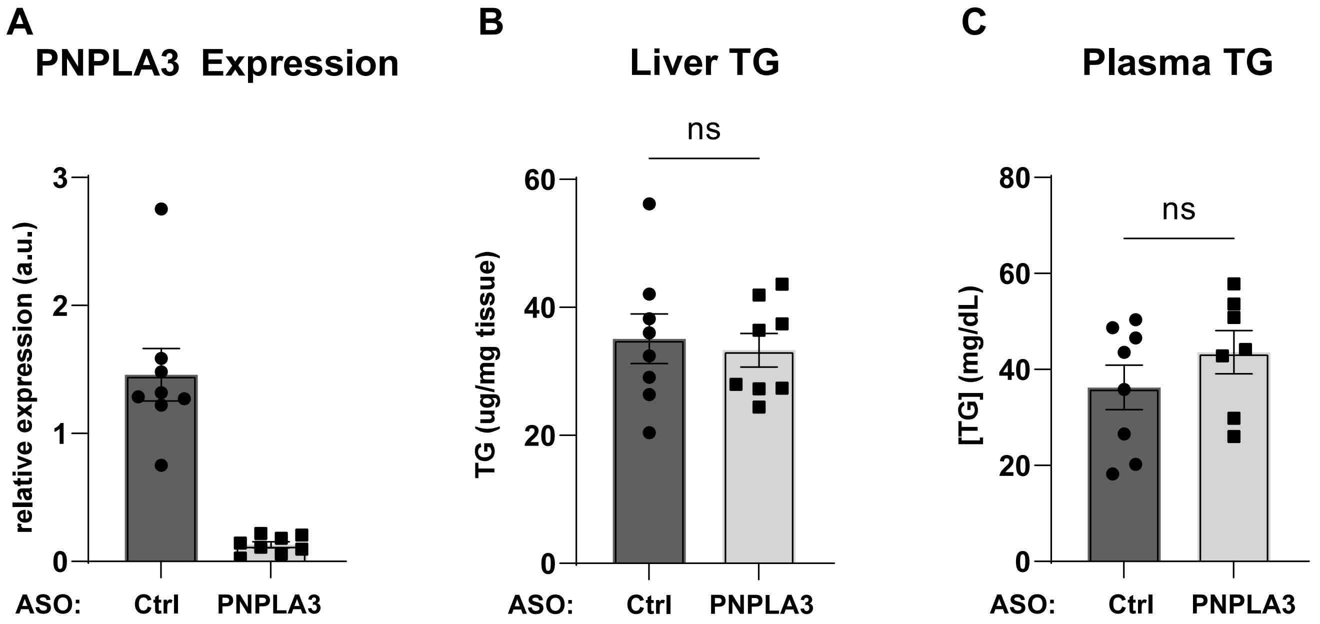

### Supplemental Figure S4

Supplemental Figure S4

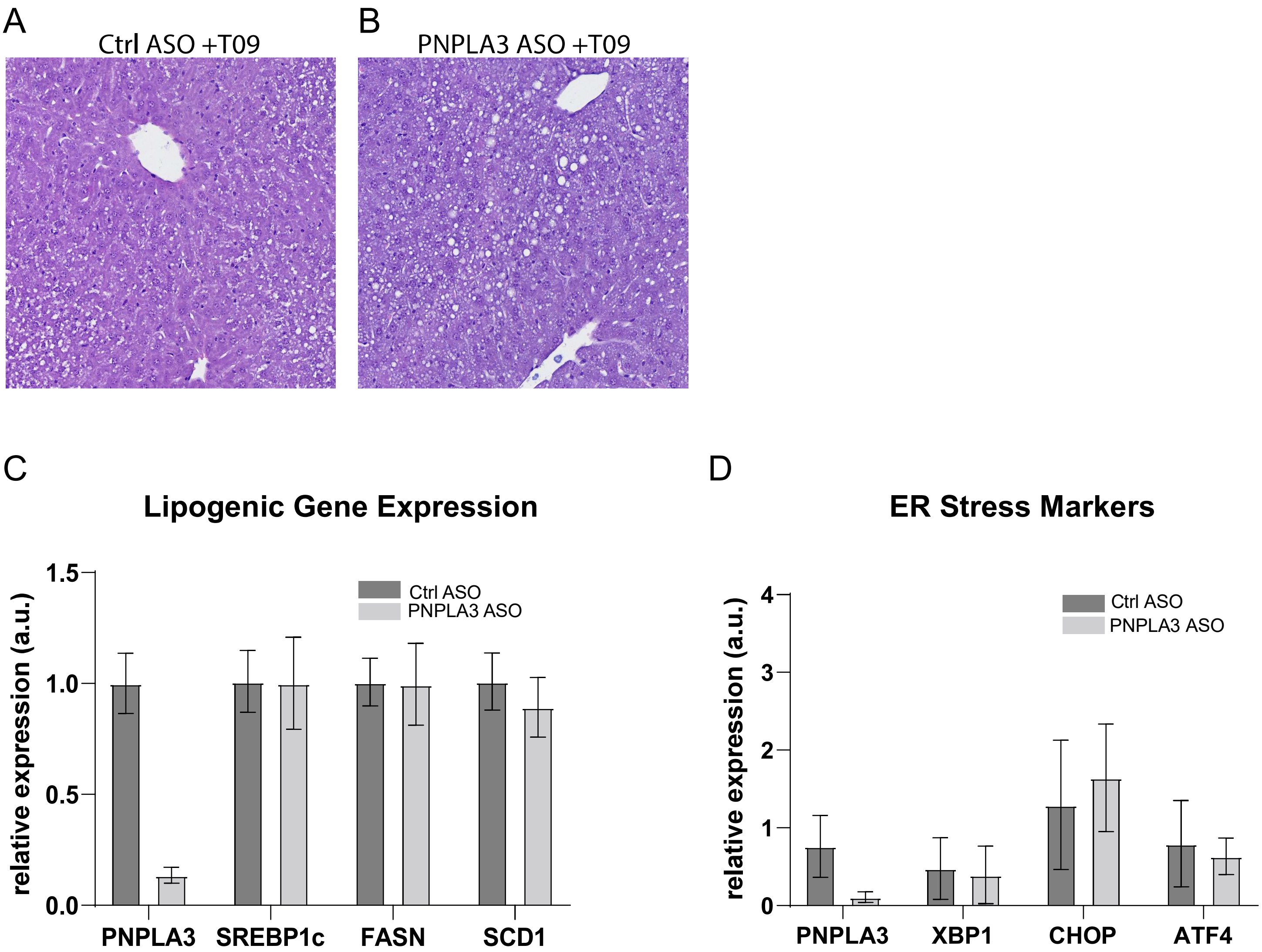

### Supplemental Figure S5

#### Supplemental Figure S5

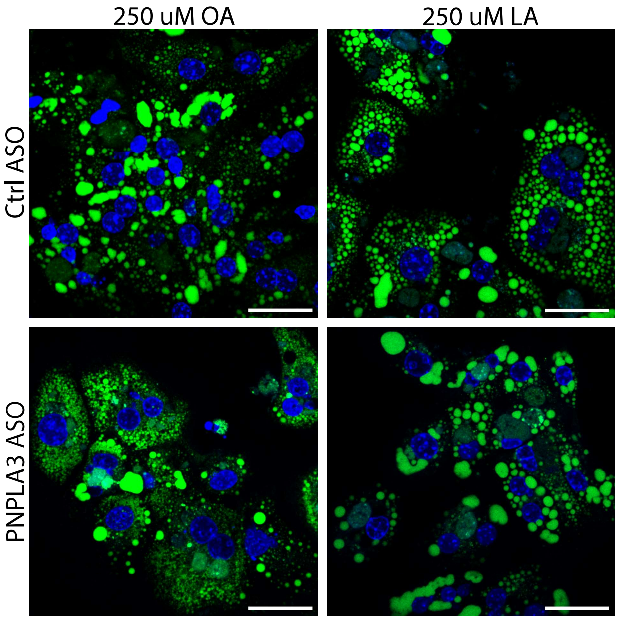
